# Ontology-based similarity calculations with an improved annotation model

**DOI:** 10.1101/199554

**Authors:** Sebastian Köhler

**Affiliations:** NeuroCure Cluster of Excellence, Charité Universitätsklinikum, Charitéplatz 1, 10117 Berlin, Germany

**Keywords:** ontologies, phenotype, gene ontology, semantic similarity, study-wise annotations

## Abstract

A typical use case of ontologies is the calculation of similarity scores between items that are annotated with classes of the ontology. For example, in differential diagnostics and disease gene prioritisation, the Human Phenotype Ontology (HPO) is often used to compare a query phenotype profile against gold-standard phenotype profiles of diseases or genes. The latter have long been constructed as flat lists of ontology classes, which, as we show in this work, can be improved by exploiting existing structure and information in annotation datasets or full text disease descriptions.

We derive a study-wise annotation model of diseases and genes and show that this can improve the performance of semantic similarity measures. Inferred weights of individual annotations are one reason for this improvement, but more importantly using the study-wise structure further boosts the results of the algorithms according to precision-recall analyses. We test the study-wise annotation model for diseases annotated with classes from the HPO and for genes annotated with Gene Ontology (GO) classes. We incorporate this annotation model into similarity algorithms and show how this leads to improved performance.

This work adds weight to the need for enhancing simple list-based representations of disease or gene annotations. We show how study-wise annotations can be automatically derived from full text summaries of disease descriptions and from the annotation data provided by the GO Consortium and how semantic similarity measure can utilise this extended annotation model.

## Background

Ontologies have become a widely used tool to capture knowledge about objects in biology, genomics, and medicine. Besides enabling knowledge integration and retrieval, they are also a widely used tool for similarity calculation between items that have been described (annotated) with classes of an ontology (1). Ontology-based similarity measures allow non-perfect matches between ontology-classes to be quantified by incorporating the graph-structure of the ontology. Often used similarity measures include semantic similarity measures based on Resnik’s definition of shared information content (2), cosine similarity measure, and the Jaccard index (1).

In the fields of human genetics, genomics, and precision medicine, the Human Phenotype Ontology (HPO, (3,4)) is often chosen to store the information about the clinical features of patients in a computer-interpretable way. In most applications of the HPO, a similarity measure is used to compare a phenotype profile of a patient against a set of diseases or genes, which itself are represented as a gold-standard set of HPO classes (5–7). The result of this step is a quantification of the similarity or overlap of the query profile and the gold-standard profile. Similarly, a typical application of the Gene Ontology (GO, (8)) is the search for similar proteins for a given query protein described as a set of GO classes known to be associated with this protein.

A critical part of this process is the comparison of a query set of ontology classes against a gold-standard set of ontology classes associated with an item, such as a disease or a protein. These gold-standard profiles are often represented as flat lists.

In human genetics, gold-standard profiles were constructed from human-written summary tables of clinical features. For example, the clinical synopsis section of the Online Mendelian Inheritance in Man database (OMIM, (9,10)) database provides a tabular view of the clinical features seen in patients with a particular disease. Orphanet provides a similar list of clinical features for each Orphanet disease entry, but encodes these directly as HPO classes (11,12). Accordingly, the HPO project, until now provides flat lists of HPO classes for each disease or disease gene, which we call “Merged” annotation model.

However, full text disease descriptions often have an inherent structure, which we aim to use in this work. In OMIM’s full text description, each paragraph often represents a summary of a certain publication, study, or disease aspect. Similarly, GO-annotations often have references to PubMed articles where a particular annotation has been derived from.

The hypothesis of this work is that standard semantic similarity measures can be improved by extending the standard “Merged” annotation model with additional information (“By study” annotation model) that can be easily extracted from the accompanying full text description in OMIM and the meta-information available in the GO annotation data set.

## Methods

We first show how we construct a study-wise annotation model for annotations with the Human Phenotype Ontology (HPO) and Gene Ontology (GO). We introduce semantic similarity measures that take into account weights of individual phenotypes and the study-wise structure of the annotation data. Finally, we present the data and approach we chose for evaluating the hypotheses of this work.

### Text mining of OMIM full text

We obtained one of the last freely downloadable versions of omim.txt (last updated on January 8th, 2016) and wrote a parser that extracts the full text description for all OMIM entries that the HPO project provides annotations for (6916 in total). The full text description of OMIM is structured according to multiple sections, such as “biochemical features”, “molecular genetics”, or “clinical features”. We aimed to identify HPO classes associated with patients that are diagnosed with OMIM entries. To prevent false positive associations, we restrict our text mining procedure to the sections “description”, “other features”, “biochemical features”, “diagnosis”, “clinical features”, and the introductory section that usually has no header.

The text in the OMIM-sections is organised into multiple paragraphs - it is this paragraph structure that we aim to investigate in this work. To give an example, the OMIM entry for Alzheimer disease (OMIM:104300) contains the paragraph

> *Yan et al. (1996) reported that the … was particularly increased in neurons close to deposits of amyloid beta peptide and to **neurofibrillary tangles***.

This paragraph is later followed by the paragraph

> *Bergeron et al. (1987) found that … findings suggested that **cerebral amyloid angiopathy** is an integral component of AD*.

The assumption is that each paragraph roughly corresponds to one study. We thus splitted the text into separate paragraphs and used the NCBO annotator (13) to identify HPO classes in each paragraph. We then stored the found matches together with the paragraph index, i.e. we generate a file that contains lines such as

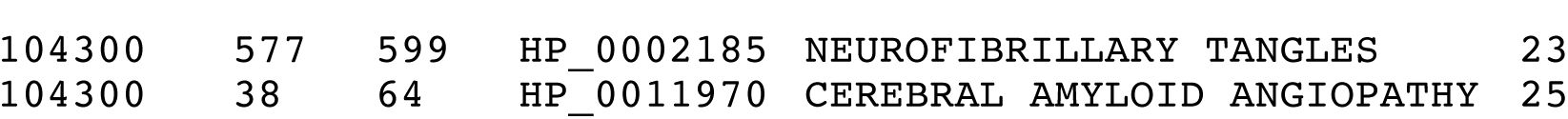

which means that NCBO annotator found the HPO class for “Neurofibrillary tangles” (HP_0002185) in the 23^rd^ paragraph of the OMIM entry with the ID 104300. It also lists the indices of the region spanned by the match (577 to 599). The next line lists the occurrences of “Cerebral amyloid angiopathy” (HP_0011970) in the 25^th^ paragraph of the same OMIM entry. We will infer from this data the study-wise annotation model, by assuming that each paragraph represents one study.

### Gene Ontology Annotations

We downloaded the GO obo-version from http://purl.obolibrary.org/obo/go-basic.obo (date: 2017-05-10) and the GO annotation data from ftp://ftp.ncbi.nlm.nih.gov/gene/DATA/gene2go.gz (date: 2017-05-10). Note that we only considered GO classes from the biological process (BP) sub ontology. We used all human *gene to GO-BP* class associations that have at least one PubMed reference, as we used those for deriving the study-wise annotation sets for each gene.

### Annotation models

In this work, we test four different annotation models which can be generated from the annotation data described before. We introduce a merged (“Merged”), weighted (“Merged (weighted)”), study-wise (“By study”), and randomised study-wise model (“By study (shuffled)”). An illustration of these models is shown in Figure 1.

**FIGURE 1.**
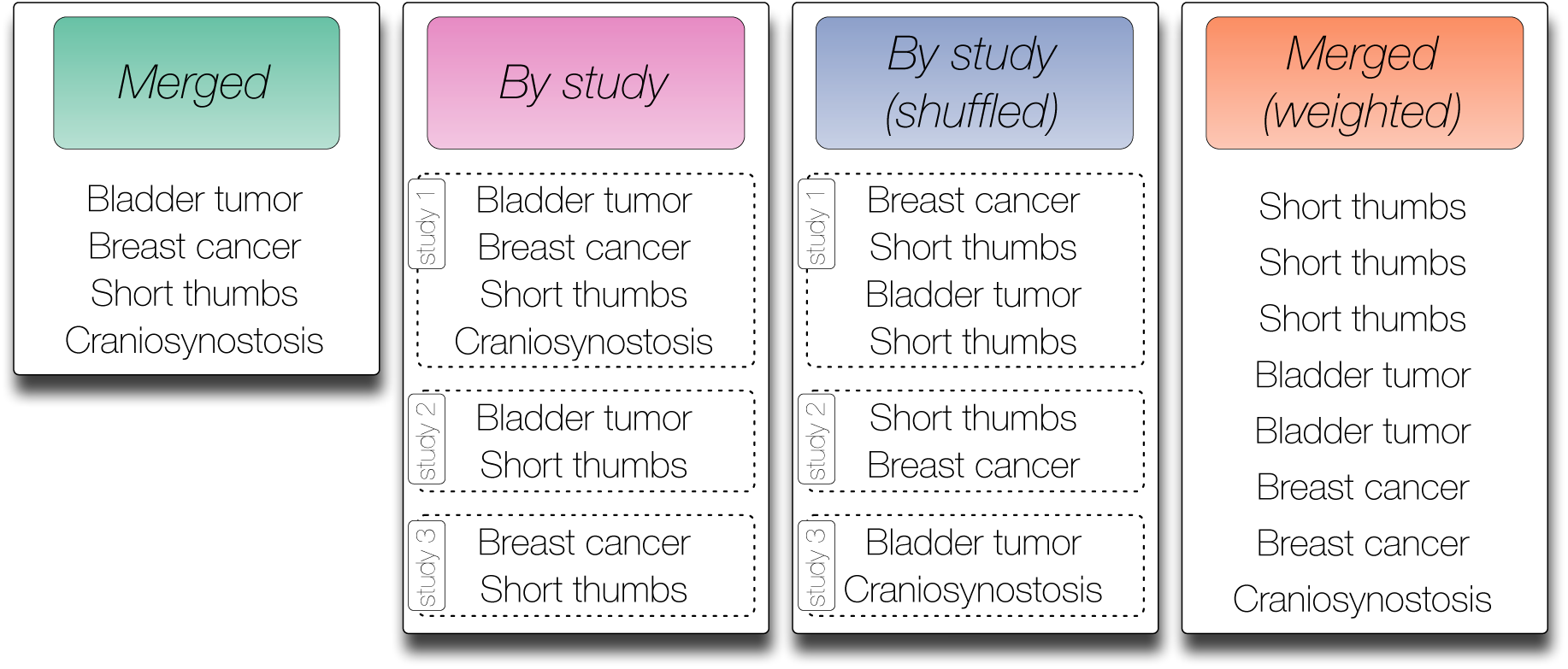
Contrived symptom annotation set for one disease under the different annotation models tested in this study. In the “Merged” model every symptom is listed exactly once and no further structure is given. In the “By study” model the symptoms associated with the disease are organised by three studies that mention the symptoms. Note that each symptom may be mentioned in multiple studies. In the shuffled version of the “By study” model each symptom from a study is randomly switched with a symptom from another study. Finally, the “Merged (weighted)” model removes the structure defined by the studies but keeps the number of mentions for each symptom as derived from the “By study” model.

The merged annotation model (“Merged”) resembles the model that is currently widely used, e.g. in the annotation file provided by the HPO. Here, each ontology class is listed once in a flat list with no additional structure. The study-wise annotation model (called “By study”) creates an annotation list for each study. For the HPO this corresponds to each paragraph of the full text disease description from OMIM that was considered in this work (i.e. we did not consider all sections as described before). For GO this corresponds to each PubMed article referenced in the annotation data.

To investigate the influence of the actual annotations per study and to exclude other factors introduced by the studies (e.g. the number of studies per disease/gene), we also use a randomised version of this annotation model (called “By study (shuffled)”), where we kept the number of studies and the number HPO/GO classes per study constant, but randomly exchanged annotations between studies of each disease/gene. We perform this exchange between two randomly chosen HPO/GO classes from two randomly chosen studies 1000 times per disease/gene.

One of the obvious differences between the annotation models “Merged” and “By study” is the number of associated HPO/GO classes per study, because multiple occurrences of one ontology class per disease/gene are allowed in the latter model (see Figure 1). To investigate the influence of this criterion, we also introduce the annotation model “Merged (weighted)”, where we use one flat list of associated ontology classes, but each class is listed as often as it occurs in different studies. Figure 1 illustrates the four different annotation models using an artificial example disease and artificial HPO annotations.

### Similarity measures

We chose to test three widely used semantic similarity measures to quantify the overlap between a query Q and an item I (an item is a disease or a gene in our case). Here, each of these measures is capable to take into account the number of occurrences of a particular HPO/GO class in Q and in I. This means, that measures are able to make use of duplicated annotations in the model “Merged (weighted)”. We also introduce a method to apply the measures to the study-wise annotation model.

#### Unweighted and weighted semantic similarity measures

We took a standard semantic similarity measure, which is based on Resnik’s definition of information content (IC). For each class c in the ontology, the information content IC(c) is defined as the negative logarithm of the frequency of annotations of items with the class (2), i.e. IC(c) = − log(p_c_), where p_c_ is the observed frequency of items annotated with class c among all annotated items.

The similarity between two ontology classes is then calculated as the IC of their most informative common ancestor (MICA), i.e. the common ancestor with the highest IC (2). For this paper, we define the semantic similarity between the annotated ontology classes of a query (Q) and the annotated classes of an item (I) as

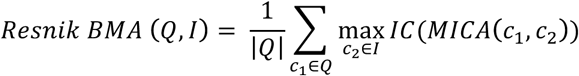

Note that |Q| returns the number of ontology classes in the set Q. For the unweighted version, each ontology class in Q and I is uniquely present, for the weighted version the sets Q and I may contain duplicate entries. BMA stands for best match average (1).

Other widely used similarity measures are the Cosine similarity and the Jaccard index. To apply these, the query Q is transferred to a vector representation q, where each entry represents the number of occurrences of the corresponding ontology class. For the unweighted version, each of the entries is set to 1, if the corresponding ontology class is present in the set and 0 otherwise. We do the same for the item I, i.e. it is transferred to the vector i. Note that during the transformation, all the ancestors of the classes in Q and I are set to the corresponding count (or 1 in the unweighted case).

The Cosine similarity measures the cosine of the angle between two non-zero vectors (here q and i) and is defined as

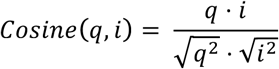

Finally, we tested the Jaccard index. The input is the same as for the Cosine measures, i.e. the transformation of Q and I into a vector representation q and i. The Jaccard index is then computed as

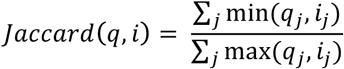

Again, in the unweighted setting, each entry in the vectors can be either 1 or 0, but in the weighted setting the entries represent the number of occurrences of the corresponding ontology class in Q (or I).

#### Paragraph-wise similarity measures

In this work, we extend the flat list representation of gold standard annotations (Q, I) by a study-wise model, such that we replace the list of annotated HPO or GO classes per disease or gene with multiple separate lists per disease or gene (see Figure 1). The similarity in the study-wise annotation model is the defined as

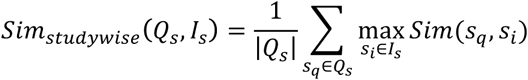

where Q_s_ and I_s_ are defined as the set of studies Each study s_x_ represents a list of ontology classes that were identified in the corresponding study. This simply means that for each study in the query, we try to identify the most similar study for the item. The final score is just the average of similarities between the studies. The function *Sim(p_q_, p_d_)* is a placeholder for one of the standard semantic similarity measures described in the previous section, i.e. Resnik BMA, Cosine, or Jaccard. Intuitively, this method takes each paragraph from the query and tries to identify the best matching paragraph in the item.

### Evaluation using OMIM phenotypic series and pathway membership

In order to test the different annotation models, we chose to use pre-defined groupings of diseases and genes, where the goal is to re-identify group members given one member as query.

#### HPO-phenoseries test

For HPO we used OMIM phenotypic series data, which is a tabular representation for viewing at genetic heterogeneity of similar phenotypes (9). One example of a phenotypic series is “Reticulate pigment disorders”, which has six members. The idea is to use the HPO annotations of one member of a phenotypic series as query (e.g. “Dowling-Degos disease 1”, OMIM:179850) and rank all OMIM entries by semantic similarity to that query. We record the rank of the other members of the phenotypic series (e.g. “Dyschromatosis symmetrica hereditaria”, OMIM:127400), aiming to rank the other members as highly as possible, i.e. above all non-members. The test only considered phenoseries groups with at least 2 diseases after removing all diseases that had less than 5 HPO annotations and less than 4 different studies. This applied to 233 phenoseries.

Please note that we did not aim to generate a more detailed or deeper HPO annotation data set than the existing one, but rather want to show that for the one annotation set derived from mining OMIM full text descriptions, meta information on the annotations contains valuable information. We thus did not compare the performance to the existing gold-standard flat list: The aim was to show that for this annotation set, a more sophisticated model such as the “By study”-annotation model outperforms a flat list (“Merged”) representation which is currently applied.

#### GO-BP-pathway test

For GO, we obtained gene-to-pathway associations from KEGG (14), by downloading the file http://rest.kegg.jp/link/pathway/hsa (date: 2017-09-19). Similar to the OMIM test, we took one gene from each pathway and used its GO annotations to the “biological process” sub ontology (GO-BP) to calculate a similarity to all other genes. The test only considered pathways with at least 2 genes but not more than 15 genes left after removing all genes that had less than 5 GO-BP annotations and less than 4 different studies. This applied to 89 pathways.

#### Performance evaluation

As written before, we test the performance of the different similarity measures and annotation models by trying to identify all group members given one member as query. We record the ranks of all other members; whereby low ranks are better. We draw the rank distribution of the obtained ranks and plot them as box- and violin plots. A boxplot shows 50% of the data points surrounding the median in the box and the position displays the skewness of the data. The whiskers extend to the most extreme data point that is no more than 1.5 times the length of the box away from the box. Violin plots are similar, except that they also show the probability density of the data at different values. We use an overlay of a box- and violin-plot. To test for significant improvements of the rank distributions, we use Wilcoxon signed rank test with continuity correction, a non-parametric test for two unrelated, not normally distributed samples (function *wilcox.test(x,y,paired=T,alternative=”less”)* in R).

We also use precision/recall curves (PRC), which are visual representations of the performance of a model in terms of the precision and recall statistics. For different thresholds, it plots the actual precision (y-axis) and recall (x-axis) points and connects them by a line. An important measure is the area under this curve (AU-PRC), which is to be maximised.

## Results

In this work, we tested four different annotation models, in particular we investigate a model that incorporate the information about the study or publication that a particular annotation has been based upon (see Methods). Figure 1 illustrates the different annotation models tested in this work.

### GO and HPO annotation data sets

In total, we annotated 6475 OMIM entries with at least one HPO class from the HPO sub ontology “Phenotypic abnormality” (HP:0000118) using the NCBO Annotator (13). For these OMIM entries we have identified 29,202 paragraphs/studies with at least one HPO class, i.e. on average each disease has 4.5 paragraphs/studies in our dataset. In the “Merged” annotation model each disease has on average 14.1 HPO annotations, whereas in the “By study” annotation model, each disease has 21.3 HPO annotations. The reason for this discrepancy is the missing uniqueness constraint. Each study/paragraph has a mean number of 4.73 HPO annotations.

For the GO test we used 10,608 genes that are annotated with at least one GO class from GO-BP (biological process). We have identified 30,104 PubMed-references and on average each gene has 2.8 PubMed-references (studies). Using the “Merged” annotation model each gene has on average 4.5 GO-BP annotations. In the “By study” annotation model, each disease has 5.1 GO-BP annotations. Each study/PubMed-reference has a mean number of 1.8 GO-BP annotations.

The distribution of the annotation size (i.e. the distribution of the number of ontology classes associated with an item or a study) is here analysed using quantiles (function quantile in R). Here we report the numbers for which five 5% and 95% of the data is smaller. For all tested annotation models (GO-BP “Merged”, GO-BP “By study”, HPO “Merged”, HPO “By study”) the 5% quantile is 1. However, the 95% quantile differs significantly between the “Merged” and the “By study” annotation data set, i.e. 15 vs. 4 for GO-BP and 43 vs. 14 for HPO). This shows that the annotation size distribution for the “By study” model is more homogeneous than the distribution for the “Merged” model.

### Performance of study-wise similarity measures

We analysed if the annotation model currently in use can be improved by employing a study-wise annotation model. For the Human Phenotype Ontology (HPO) we derived the study-wise annotations by analysis of OMIMs full text descriptions. For Gene Ontology (GO) we used the PubMed-identifiers often available for associations between a gene and a GO class.

We tested four different annotation models and three different similarity measures in two tests, the HPO-phenoseries and the GO-BP-pathway test (see Methods). We found that the study-wise model outperforms almost all the other models in the tests according to the modes of evaluation chosen.

Figure 2 plots the rank distributions of the sought items in the HPO-phenoseries test (2 A) and for the GO-BP-pathway test (2 B) using violin- and boxplot (see Methods). The advantage of the study-wise annotation model is especially apparent in the HPO-phenoseries test.

**FIGURE 2.**
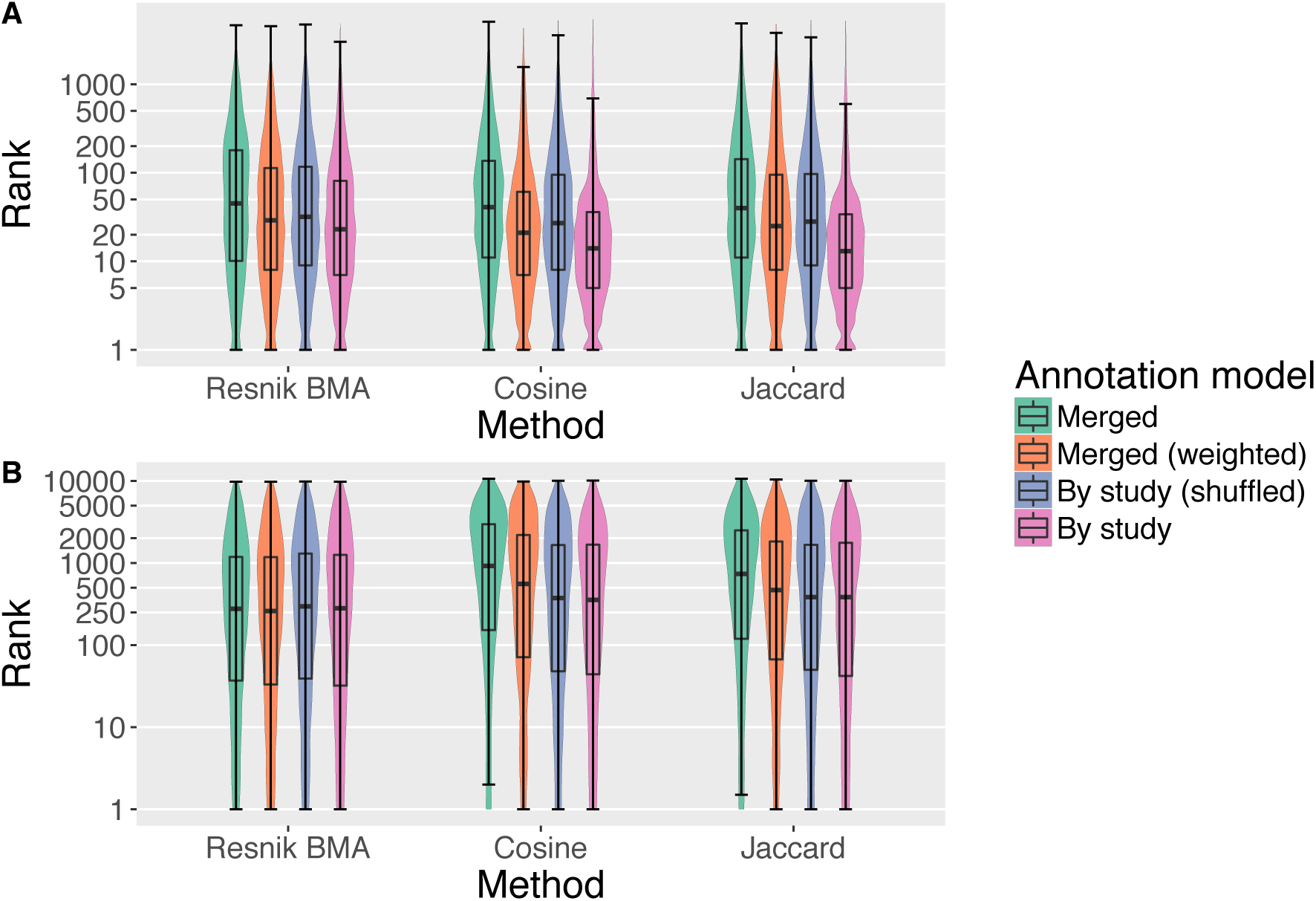
Box- and Violin plots (log-scaled) for the two tests performed in this project. A) shows the results for the HPO-phenoseries test and B) shows the same for GO-BP-pathway test. We tested three different ranking methods (Resnik BMA, Cosine, and Jaccard) and four different annotation models. The plots show the distribution of the ranks of the sought members of the corresponding group (i.e. Phenoseries members and KEGG pathway members).

Interestingly, the “Merged (weighted)” model already improves the performance of the similarity measures, but the study-wise model often outperforms the other models by a large margin. This was confirmed by Wilcoxon signed rank test (see Methods), which resulted in significant p-values for all measures in the HPO-phenoseries test, when comparing ranks using the “By study” annotation model with all other annotation models (p < 2.6*e^-161^). For the GO-BP-pathway test the results are different. Here, only the Cosine and Jaccard results are highly significant (p < 1.05*e^-128^) when comparing the merged and the study-wise annotation model. For Resnik BMA only the comparison of the ranks obtained by study-wise and the randomised study-wise model are significant (p<0.0002). For the cosine similarity measure, we also see a significant difference between the study-wise and the “Merged (weighted)” model.

In Table 1 we list the improvements in terms of increase of the area under the precision recall curve (AU-PRC). Figure 3 shows the precision recall curves for the “Merged” and “By study” annotation model for all tested similarity measures. Except for one case, the latter annotation model obviously improves the results of the precision recall analysis. Only the GO-BP pathway test for the Resnik methods show almost no change with only a negligible increase of the AU-PRC of 0.007 (c.f. Table 1). In Figure 4 we see the precision recall curves for the Jaccard similarity measure, which again shows that the “Merged (weighted)” and “By study (shuffled)” models show an improvement. As seen before, the study-wise structure contains valuable information which leads to a further performance improvement.

**TABLE 1.** Improvement by using the “By study” annotation model in comparison to the “Merged” annotation model measured by area under the precision recall curve (AU-PRC).

| ALGORITHM | INCREASE AU-PRC WITH “BY STUDY”<br>ANNOTATION MODEL |  |
| --- | --- | --- |
|  | HPO-Phenoseries<br>test | GO-BP-Pathway<br>test |
| RESNIK BMA | 0.07 | 0.006 |
| COSINE | 0.19 | 0.028 |
| JACCARD | 0.20 | 0.027 |

**FIGURE 3.**
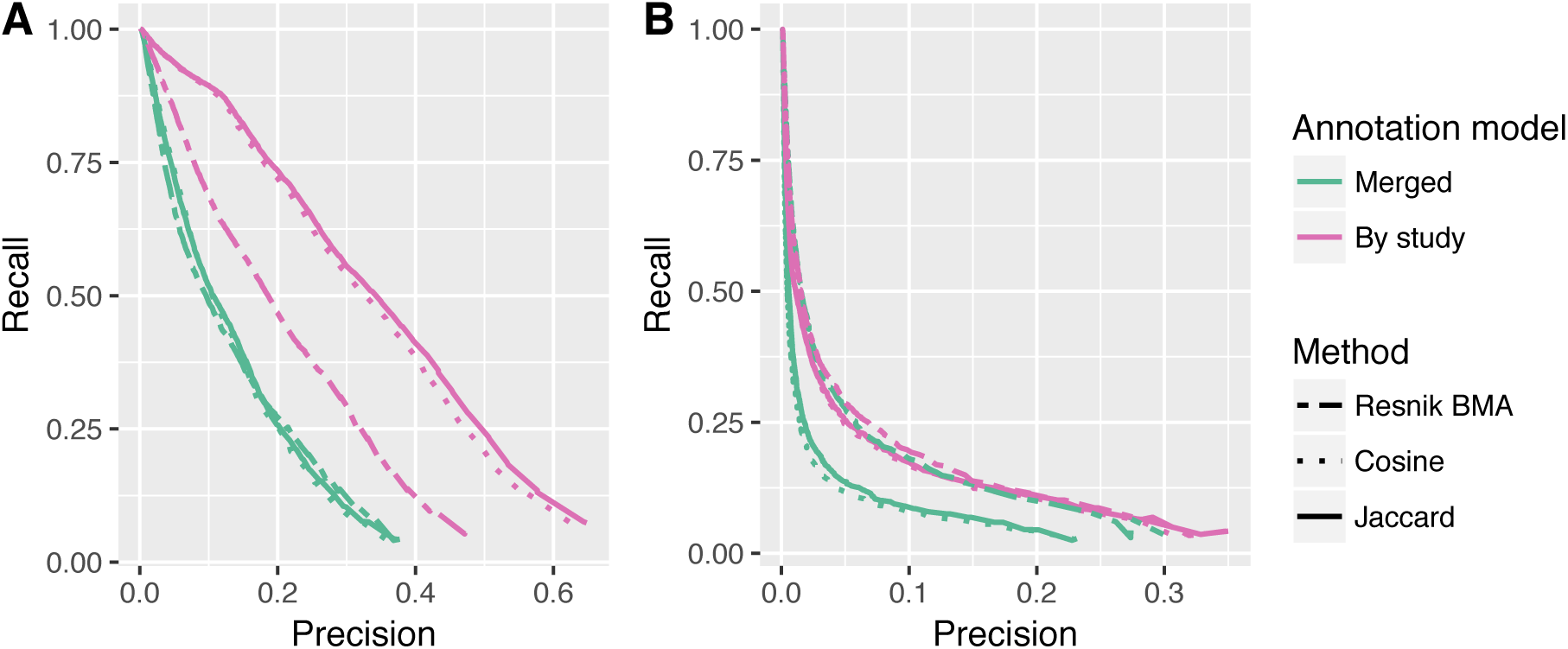
Precision recall curves (PRC) for two annotation models (“Merged” and “By study”) and for the three different ranking methods. A) and B) show the results for the HPO-phenoseries test and the GO-BP-pathway test, respectively. The differences in the area under PRC are listed in Table 1.

**FIGURE 4.**
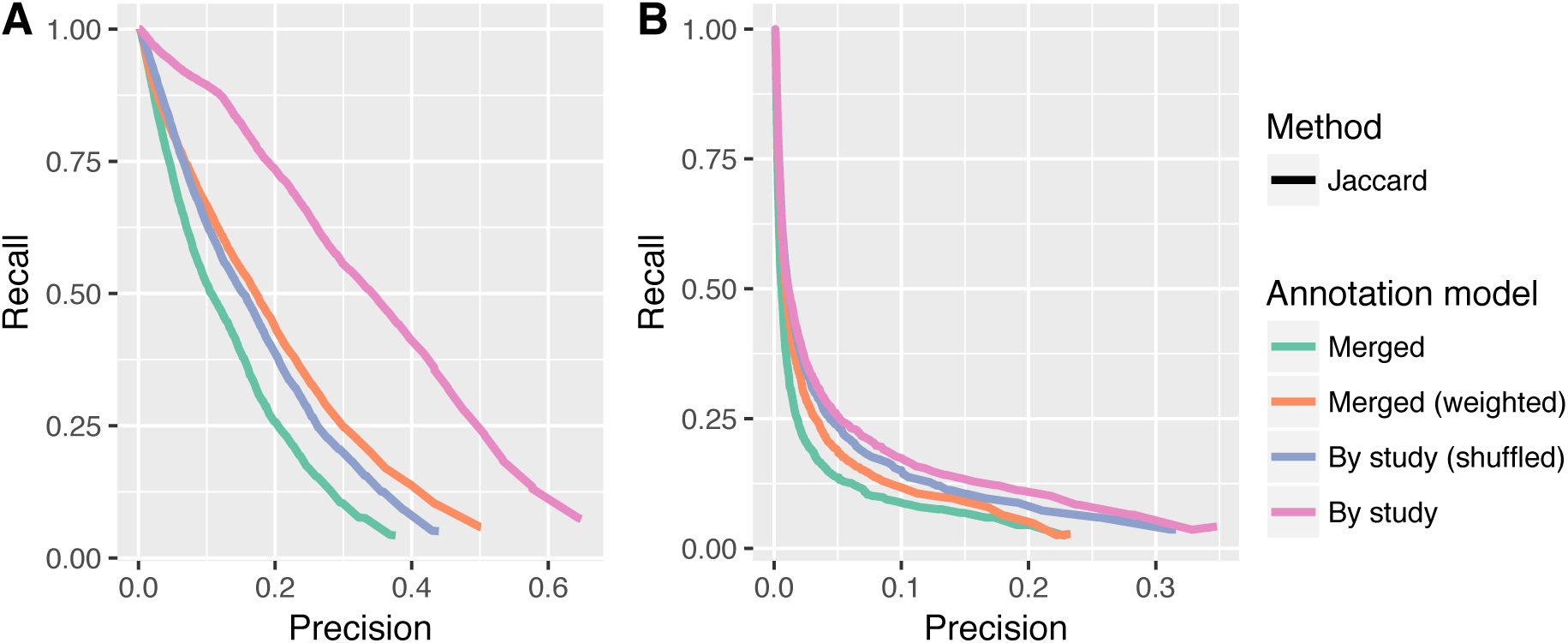
Precision recall plots for the Jaccard method and all four annotation models tested in this work. A) and B) show the results for the HPO-Phenoseries- and GO-BP-pathway test, respectively. One can see that the “Merged (weighted)” model already improves the performance, but that the study-wise model outperforms all other models.

## Discussion

This work has investigated if using one simple flat list of ontology classes for describing an item is an optimal annotation model. Usually multiple studies are underlying such an annotation set and we have thus inferred a study-wise annotation model for disease being annotated with classes of the HPO and for genes being annotated with classes of the GO-BP. We tested different semantic similarity measures that are capable of using both annotation modes and analysed their performance in recovering item-groupings defined by biomedical knowledge, i.e. OMIM phenoseries members using HPO and KEGG pathway members using GO-BP. We find that the study wise annotation model in almost all tests significantly outperforms the merged annotation model. The results show that the hypothesis of this work is supported stronger when using HPO-phenoseries test. The difference in performance advantage probably due to the low number of annotation per study (1.8 vs. 4.7) in the GO-BP-pathway test. Also, we expect more heterogeneity in GO-BP annotations for genes in one pathway compared to rather homogeneous HPO annotations for members of a phenotypic series.

The HPO-phenoseries test investigated the ability of the presented ideas to improve disease-clustering. However, often patient phenotypes are compared against disease entities and it will be interesting to see if the ideas presented here also improve performance of similarity algorithms in this setting (15), even if it is impossible to generate study-wise annotations for a single patient.

The results obtained by the study-wise approach are not only important for developers of ontology based algorithms for semantic similarity but also an important message for database curators and developers. It is essential to keep the provenance of annotations, i.e. why a particular association between an item and an ontology class has been made. Unfortunately, this has not been done for the “clinical synopsis” in OMIM (at least not publicly available) and will require a significant amount of work in the future to add this information. However, it is relatively simple for database curators to keep track of the provenance of associations in the future.

More research into the underlying mechanism of the demonstrated improvements is needed. One explanation might be the exclusion of false combinations of annotations in the study-wise annotation model, which are otherwise introduced by merging all annotated ontology classes into one list. Another explanation might be that by using study-wise annotations sets, we derive a way more homogeneous distribution of annotation sizes (see Results) with less entries having an extremely high number of annotations. The effect of annotation size on the performance of semantic similarity measures has been subject of research in recent years (15–17).

Future work will include the implementation of the presented ideas into Bayesian algorithms (18). We think the results of this work put weight on the need for improved annotation data for items in parallel to continued research into semantic similarity algorithms. This work is not only relevant for HPO and GO-BP, but also for all ontologies that are used to describe the properties and characteristics of items using ontologies.

